# Single-cell analysis of brain-derived *Toxoplasma* bradyzoites reveals a novel cell cycle regulated by AP2XI-6

**DOI:** 10.1101/2025.10.20.683543

**Authors:** Emma Franklin, Argenis Arriojas Maldonado, Kyra Lee, Grace Hilliard, Bruno Martorelli Di Genova, Gary E Ward, Kourosh Zarringhalam, Robyn Kent

**Affiliations:** Department of Microbiology and Immunology, University of Oklahoma Health Science, Oklahoma City, 73104, OK, United States; Center for Personalized Cancer Therapy, University of Massachusetts Boston, Boston, MA, 02125, United States; Dept of Microbiology and Molecular Genetics, University of Vermont, Burlington, VT, 05405, United States; Department of Mathematics, University of Massachusetts Boston, Boston, MA, 02125, United States

**Keywords:** *Toxoplasma gondii*, tissue cysts, cell cycle, chronic infection, bradyzoite replication, bradyzoite heterogeneity

## Abstract

*Toxoplasma gondii* prevalence is due, in part, to its ability to persist in hosts while retaining the capacity to transmit and recrudesce. A process that is poorly understood. Through single-cell RNA profiling of *in* vivo-derived bradyzoites, we discovered that they are heterogeneous and not G1-arrested, as expected from *in vitro* studies. Instead, they progress through two cell cycles that branch from a single G1. While in G1b, *in vivo*-derived bradyzoites express cyst wall proteins predicted to replenish the wall that shields cysts. One cell cycle is common amongst tachyzoites and *in vitro-* and *in* vivo-derived bradyzoites; the other is unique to *in* vivo-derived bradyzoites (BCC). We demonstrated that AP2XI-6, expressed in G1 *in vivo,* is dispensable for tachyzoite growth and cyst formation *in vitro* but necessary for encystation *in vivo.* We propose that AP2XI-6 is a driver of replication through the BCC that is required for cyst maturation and persistence.

## Introduction

More than one-third of the human population is infected with the prevalent pathogen *Toxoplasma gondii* [1]. Following dissemination and control of the acute tachyzoite infection by the immune system, parasites differentiate into the chronic bradyzoite stage [2–4]. Regulation of this conversion has garnered much recent attention with several transcriptional regulators being identified [5–8]. Differentiation is associated with lasting changes, including assembly of the protective cyst wall [9–17], formation of starch storage granules [18–21] and switches in metabolism [18, 22–26].

Much of what we have learned about bradyzoite biology has been uncovered from studies using an *in vitro* differentiation model [3, 27]. Single cell RNA sequencing (scRNAseq) on *in* vitro-derived tachyzoites and bradyzoites demonstrated that following asynchronous differentiation bradyzoites are heterogeneous, express many canonical stage specific markers, to variable degrees, and are predominantly G1 arrested (>90%) with infrequent replication [28]. Replication frequency was also shown to have a temporal component *in vivo.* In a cyclic pattern lasting ∼8-weeks, bradyzoites undergo periods of low replication followed by periods where all bradyzoites within a cyst and majority of cysts in the host replicate [20, 21, 29]. How bradyzoites modulate their growth for persistence and recrudescence in vivo is currently unknown.

Here we extend our understanding of bradyzoite biology by performing scRNAseq on *in* vivo-derived bradyzoites. This profiling revealed several unrecognized features of bradyzoite tissue cyst assembly and cell cycle dynamics. Surprisingly, cyst wall proteins are not ubiquitously expressed, instead their expression is confined to G1b. Integration with existing *in vitro* datasets [28] revealed the presence of novel bradyzoite subpopulations and cell cycle annotation revealed that while some bradyzoites replicate through the canonical cell cycle, the majority progress through a newly identified cell cycle. As the canonical cell cycle is utilized by tachyzoites, *in vitro-derived* bradyzoites and *in* vivo-derived bradyzoites we term this the “universal cell cycle” (UCC) [30–36]. In contrast the newly identified cell cycle is restricted to *in* vivo-derived bradyzoites, and we call this the “bradyzoite cell cycle” (BCC). These two cell cycle loops are anchored at a single, shared, G1 state. This G1 is transcriptionally divergent from *in vitro* G1, and our analysis revealed *in vivo* G1-specific expression of an AP2-domain containing transcription factor, AP2XI-6. We show AP2XI-6 is the gatekeeper of entry into the BCC as depletion has no effect on UCC-directed acute stage growth or encystation *in vitro,* while *in vivo* encystation, that utilizes the BCC, is dramatically reduced. We propose that these cell cycle trajectories are both required for persistence, balancing periods of cyst growth and population expansion following cyst reseeding.

## Results

### Bradyzoites are transcriptionally heterogeneous *in vivo*

To characterize gene expression in encysted bradyzoites we profiled >6,500 parasites isolated from the brains of chronically infected mice by single cell RNA sequencing (scRNAseq) (Figure 1A). Cluster analysis [37] uncovered 6 subpopulations (Figure 1B) whose marker genes denote distinct cellular processes (Figure 1C, Supplementary Figure 1A and Supplementary Table 2). Cluster 1 and 2 markers largely encode microneme proteins with cluster 1 also containing bradyzoite-specific microneme proteins such as AMA2 and AMA4. The separation of microneme gene expression into two clusters supports previous reports that tachyzoites contain two microneme subpopulations [38].

**Figure 1.**
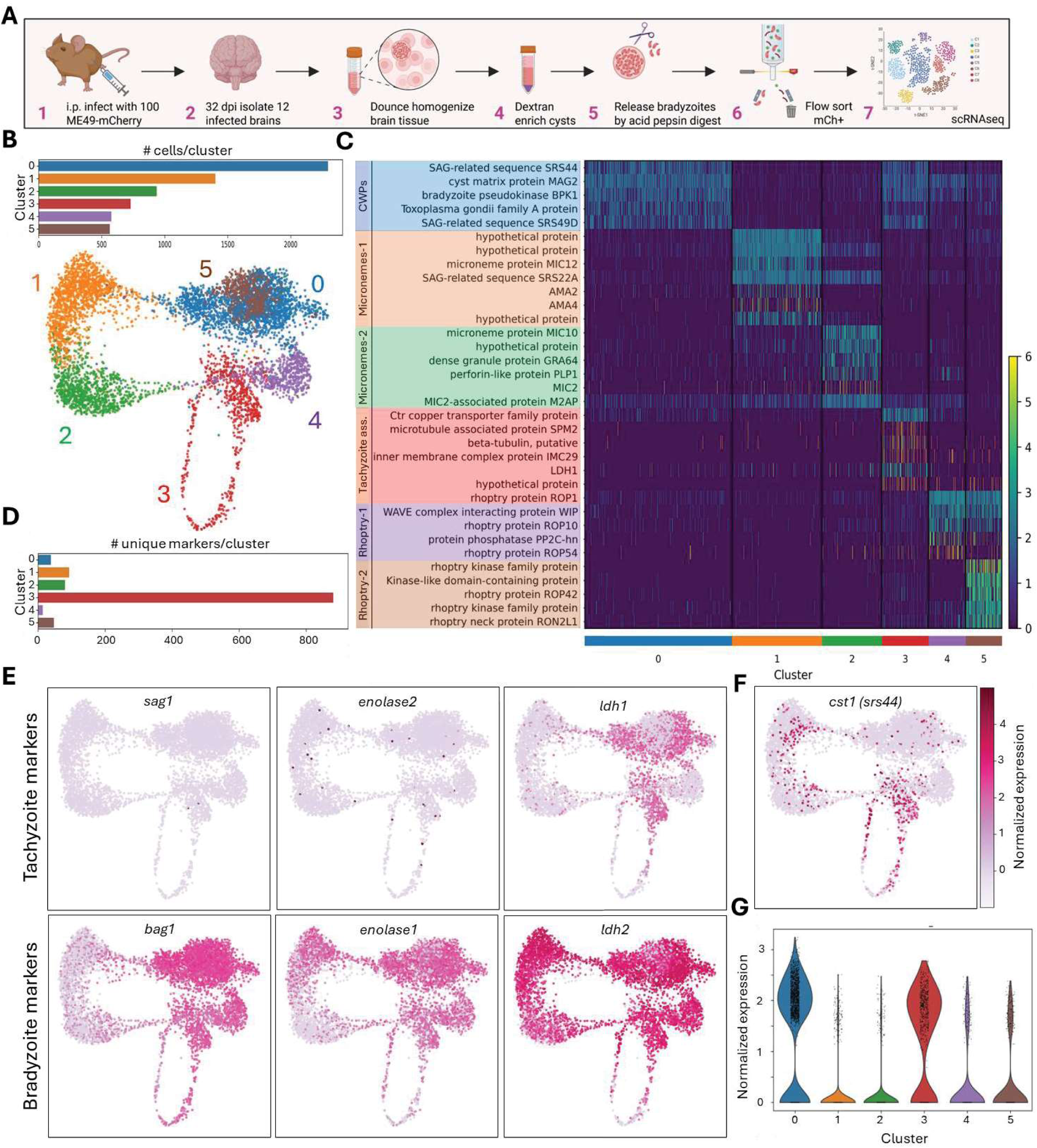
Bradyzoites are transcriptionally heterogeneous in vivo. **(A)** Schematic representation of sample preparations for single cell RNA sequencing of in vivo derived bradyzoites. **(B)** ∼6500 bradyzoites were clustered by nearest neighbor projections and the Leiden algorithm, and the data displayed as a uniform manifold approximation and projection (UMAP) to visualize clusters in 2 dimensions (bottom panel). The number of parasites in each cluster is displayed as a bar chart (top panel) **(C)** Heatmap of select marker genes for each of the 6 clusters identified by scRNAseq. Displayed markers were manually selected from all markers based on fold change compared to the whole population (log2FC > 2 and adjusted p-value < 0.05), gene annotation and function. Full heatmap and table for all markers across all clusters can be found in Supplementary Figure 1 and Table 2 respectively. From top to bottom, hypothetical proteins listed are TGME49_289370, TGME49_287040, TGME49_306270, TGME49 257970 and TGME49_ 266300. **(D)** Quantification of the total number of unique markers identified for each cluster. **(E)** UMAP of tachyzoite (top) and bradyzoite (bottom) marker genes. Individual cells are colored by normalized logCounts. **(F)** UMAP of the dominant cyst wall protein CST1 (srs44). Individual cells are colored by normalized logCounts. **(G)** Violin plot of *cst1* expression across all cluster sshows enrichment in clusters 0 and 3.

Bradyzoite specific microneme proteins are only expressed in cluster 1, along with *mic3,* suggesting only the MIC3+ microneme subpopulation contains bradyzoite specific microneme proteins after differentiation (Supplementary Figure 1B). Cluster 3 is the most divergent cluster, with >9x as many unique markers as any other cluster (Figure 1D) that includes tachyzoite associated genes (Figure 1C) [39]. Therefore, these may represent less mature, or tachyzoite-like bradyzoites within the dataset. Clusters 4 and 5 express marker genes that are predominantly annotated as rhoptry proteins (Fig 1C). Cluster 0 is enriched for expression of cyst wall proteins (CWPs) and SRS genes.

To determine if there were any de-differentiated tachyzoites in our dataset, we quantified expression of canonical tachyzoite and bradyzoite transcripts [40, 41]. Tachyzoite transcripts (Figure 1E top panels) *SAG1, ENOLASE2* and *LDH1* are infrequently (0.04%, 0.6% and 8.3%) and lowly expressed. Bradyzoite specific genes (Figure 1E bottom panels) show variable expression patterns. While *LDH2* is expressed by almost the entire population (90.7%), *BAG1* and *ENOLASE1* are only expressed by 75.7% and 68.9% respectively.

We were surprised that CWPs are markers of cluster 0 as their expression is ubiquitous in *in vitro* bradyzoites [28]. Evaluation of transcript abundance for the major cyst wall component CST1 (Figure 1F and G) and 6 other CWPs (Supplementary Figure 1C) [11, 12, 16, 17] revealed a consistent expression pattern including cluster 0 and some of cluster 3. This suggested a shared control mechanism for expression of all CWP *in vivo*.

### Spatial dynamics of cyst wall protein expression

As only a subset of bradyzoites express CWPs, we hypothesized that after initial encystation CWP expression becomes restricted to parasites at the periphery of the cyst, switching from assembly to maintenance of the wall. To test this hypothesis, we utilized RNAscope.

We fluorescently labeled the transcripts for three genes: the canonical bradyzoite transcript *ldh2* (FITC), abundantly expressed by most bradyzoites (Figure 1E), and two CWPs, *cst1* (Cy5) and *mcp4* (TRITC), expressed by 38.1% and 22% of the population respectively (Figure 1F and Supplementary Figure 1C). To filter cysts that would be useful for analysis, we quantified total fluorescence intensity to determine overall transcription levels (Figure 2A). Cysts were classified (Figure 2A and C) as having high (75%, RFU >15,000), medium (12.5%, RFU >10,000) or low (6.8%, RFU <9,000) transcriptional activity or as transcriptionally dormant (5.7%, RFU <2,500).

**Figure 2.**
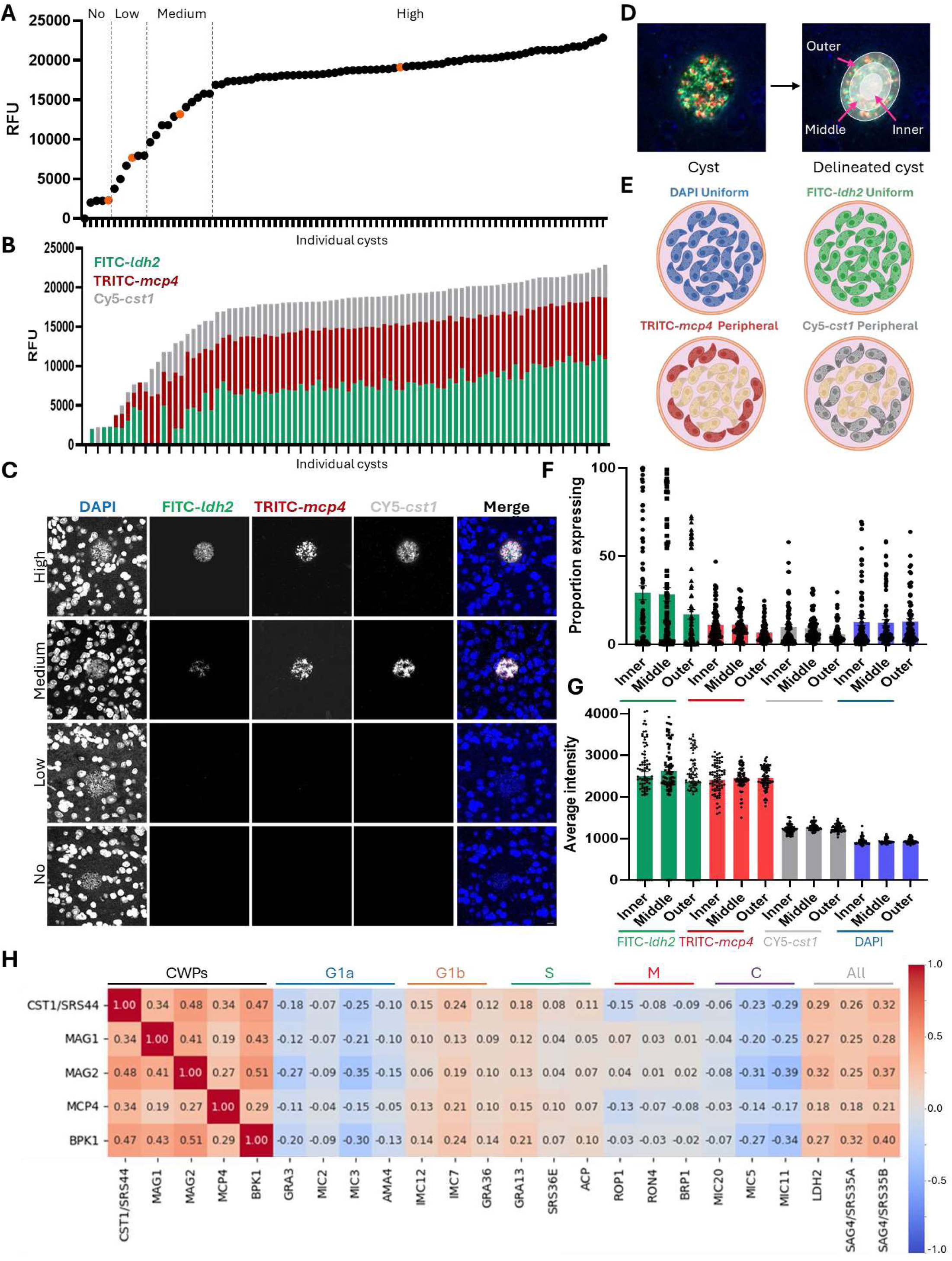
Cysts wall proteins are cell cycle but not spatially regulated. **(A)** Total (FITC-/dh2, TRITC-mcp4 and *CY5-cst1)* fluorescence intensity values for all (n= 88) analyzed cysts ordered by intensity. The data segregate into four categories: no (<2,500 RFU), low (2,501 - 9,000 RFU), medium (10,000-15,000 RFU) or high (> 15,000 RFU) expression. Orange circles are shown as representative images in C. **(B)** For each cyst the cumulative fluorescence by fluorophore is shown in sequence from lowest to highest expression. Low intensity cysts can be identified that do not express all three fluorophores, but all medium and high intensity cysts express all three transcripts. **(C)** Representative images of cysts within different fluorescence intensities shown in A. Individual fluorophores, LDH2-FITC, MCP4-TRITC and CST1-Cy5 are shown along with a nuclear stain, DAPI. **(D)** To determine if CWP expression was restricted to the periphery of cysts each of the medium and high expressing cysts (n=66) was delineated into 3 equal width concentric rings. **(E)** Model of transcript expression if CWPs are spatially regulated. Nuclear DAPI and ubiquitously expressed FITC-/dh2 will show little variability across the cyst while TRITC-mcp4 and *Cy5-cst1* expression will be limited to peripheral parasites. **(F)** Proportion of the pixels with the three regions that express each of the fluorophores. **(G)** Average pixel intensity for positive pixels within each region. **(H)** To identify any transcripts that are co-expressed with cyst wall proteins, the expression correlation coefficient between cyst wall proteins and proteins with 3 known gene that show cell cycle specific expression was calculated for each cell cycle state (Supplementary Figure 2) along with the ubiquitously expressed bradyzoite genes *sag4.* Correlation is colored to indicate a high degree of expression correlation (red) to a low correlation (blue).

As quantification of all probes (FITC-/dh2, TRITC-mcp4 and *CY5-cst1)* is optimal to determine if CWP is expression is spatially regulated, we quantified fluorescence intensity for individual transcripts. All medium and high expressing cysts showed expression of all 3 transcripts (FITC-/dh2, TRITC-mcp4, *CY5-cst1,* Figure 2B) but individual probe expression was inconsistent in the low transcriptional activity cysts. Therefore, we excluded these from our CWP analysis, leaving 66 intact cysts to be profiled.

The boundary of each cyst was manually demarked and each cyst computationally split into three equal width concentric rings (Figure 2D). If CWP expression is confined to peripheral parasites we would expect only outer-ring parasites to express TRITC-mcp4 and *CY5-cst1* while we do not expect parasite abundance (DAPI) or FITC-/dh2 abundance to vary (Figure 2E). We quantified the proportion of TRITC, FITC and CY5 positive pixels, as a measure of the number of parasites expressing each transcript in each ring and normalized by the total number of pixels in the ring (Figure 2F). We see that the outer ring contains fewer pixels with detectable level of transcription of any probed transcript. Next, we quantified the average intensity of positive pixels to determine if there was higher transcript abundance, indicating higher expression of these genes by parasites in the region (Figure 2G). This did not differ across any region for any transcript or DAPI. Taken together these data suggest that there is no difference in the number of parasites at the periphery of the cyst (based on DAPI), but the parasites at the exterior are transcriptionally less active. These data disprove our hypothesis that CWP expression is spatially regulated.

As cyst wall protein expression is not spatially regulated, we returned to our scRNAseq data to uncover the source of regulation governing co-expression of these proteins. We performed correlation analysis of 5 cyst wall proteins, with a constitutively expressed bradyzoite gene *(sag4)* and fifteen proteins with clear cell cycle dependent expression profiles in tachyzoites (Supplementary Figure 2). CWPs show co-expression with G1b expressed genes, minor co-expression with S-phase genes, and little to no co-regulation with the other cell cycle regulated transcripts (Figure 2H), showing that expression of CWPs begins when bradyzoites are in G1band continues as they transition into S-phase.

### Cysts are heterogeneous and dynamic entities *in vivo*

Interestingly, in our RNAscope data we also identified cysts that are undergoing fission, where the cyst first elongates into a kidney bean-like shape (Supplementary Figure 3A) before one (Supplementary Figure 3B) or more (Supplementary Figure 3C) cysts bud from a “parent” cyst and fully dissociate (Supplementary Figure 3D). This was previously reported *in vitro* [3] but had not been documented *in vivo.* These budding cysts account for a small subset of the population (Supplementary Figure 3E), with 4.5% forming a single bud and 1.1% forming multi-buds. We confirmed the presence of cysts undergoing fission using immunohistochemistry (Supplementary Figure 3F), where 4.6% of the cysts had a budding cyst.

### *In vivo* bradyzoites replicate via the canonical cell cycle and a novel cell cycle

Discovering the link between CWP expression and cell cycle (Figure 2H) was not wholly unexpected, as the expression of many *T. gondii* genes is governed by cell cycle progression [32]. However, previous work [28] has shown that *in vitro* >90% of bradyzoites are arrested in G1 with > 55% in G1b, whereas only 35.5% of our population express CWPs (Figure 1F and Supplementary Figure 1C) that are co-expressed with known G1b genes. This indicates that either fewer *in* vivo-derived bradyzoites are arrested in G1b or only a subset of G1b parasites express CWPs.

To determine the cell cycle state of our *in* vivo-derived bradyzoites we integrated publicly available *in vitro-derived* bradyzoite data from the ME49 strain [28] with our new *in vivo*-derived bradyzoite scRNAseq. We transferred the cell cycle annotations previously developed [28] to our *in* vivo-derived bradyzoites (Figure 3A), alongside existing data for *in* vitro-derived tachyzoites [28] (Figure 38) and *in* vitro-derived bradyzoites [28] (Figure 3C). We were surprised to discover that *in* vivo-derived bradyzoites are not G1 arrested. Instead, we identify two interconnected cell cycle trajectories with separate clusters of S, Mand C cell cycle annotated parasites (Figure 3A). While S, Mand C appear duplicated as distinct loops they both branch from a single G1a/b (blue and orange respectively, black arrow, Figure 3A). Overlaying these two datasets (Figure 3D) shows clear overlap of *in vitro-* and *in* vivo-derived bradyzoites in the lower loop where one cell cycle is traversed (Figure 3D, pink arrow). We therefore call this the universal cell cycle (UCC). Conversely, we observe in the upper loop that the second cell cycle is sparsely utilized by *in* vitro-derived bradyzoites (green arrow) leading to its designation as the bradyzoite cell cycle (BCC).

**Figure 3.**
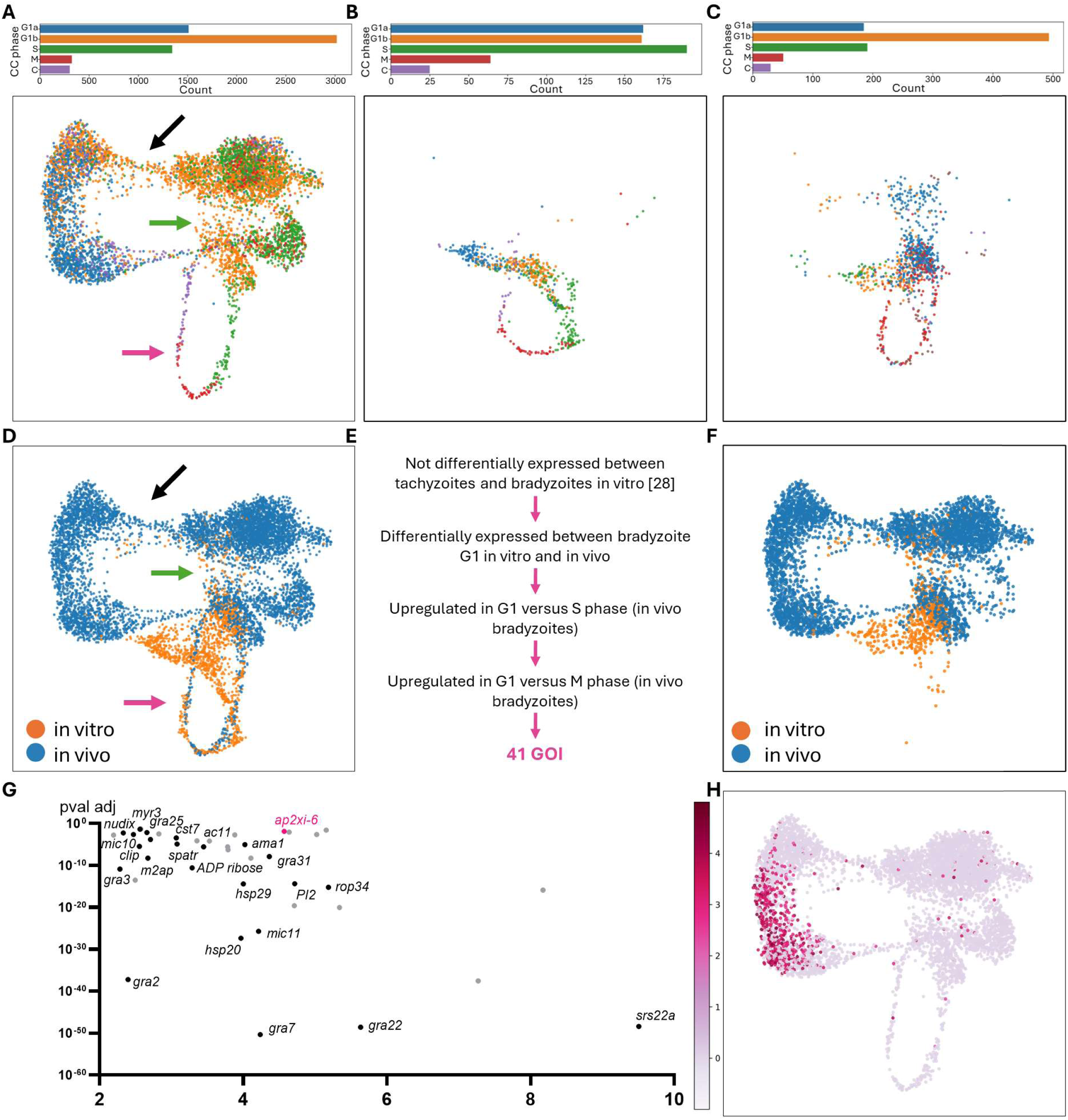
Bradyzoites can replicate through 2 concurrent cell cycles. **(A)** Cell cycle states were assigned to each of the >6,500 in vivo-derived parasites based on cell cycle state determination for in vitro tachyzoites and bradyzoites [28]. **(B)** Integration of in vitro bradyzoite and **(C)** tachyzoites datasets [28] show the presence of a single cell cycle loop. **(D)** Combining the in vitro (orange) and in vivo (blue) bradyzoite datasets we see that in vitro bradyzoites have limited overlap with in vivo bradyzoites. The lower loop (pink arrow) is common, encompassing one cell cycle now termed the universal cell cycle (UCC) while the upper loop (green arrow) is only traversed by in vivo bradyzoites and is now referred to as the bradyzoite cell cycle (BCC). These loops branch from a shared region that includes G1a and G1b (black arrow). **(E)** Flow chart of method used to identify potential BCC regulators (Supplementary Figure 5). GOI = genes of interest. **(F)** UMAP shows G1 of in vitro and in vivo bradyzoites is divergent leading us to propose that this divergence results from expression of BCC driving genes in vivo as this cell cycle is not present in vitro. **(G)** This analysis identified 41 differentially expressed genes, specifically expressed in G1 by in vivo derived bradyzoites. **(H)** The AP2 domain containing protein TGME49_215895, herein named AP2XI-6, shows clear upregulation in vivo and a G1 specific expression profile.

As *in* vivo-derived bradyzoites appeared considerably more diverse than *in* vitro-derived bradyzoites we performed cluster analysis on the combined datasets. As previously demonstrated, *in* vivo-derived bradyzoites show a great deal of heterogeneity with 6 clusters (numbered 0-5, Figure 1B and Supplementary Figure 4A). However, *in vitro*-derived bradyzoites were far less diverse, with the majority (>50%) assigned as within cluster 0 (Supplementary Figure 4B). Cluster 0 contains parasites in G1b that express CWPs in keeping with previous reports that *in* vitro-derived bradyzoites are arrested in G1 [28].

Differential expression analysis between *in vitro-* and *in* vivo-derived bradyzoites revealed upregulation of 664 and downregulation of 1443 genes *in vivo* (Log2FC > 2, p-value <0.05; Supplementary Figure 4C and Supplementary Table 3). This includes upregulation of factors known to regulate bradyzoite development (AP2IX-4 [42], AP2IV-4 [43] and BFD1 [5]), repress sexual stage development (AP2XI-2 [44]), and repress developmentally regulated tachyzoite genes (AP2XII-5 [45]), and downregulation of sexual and oocyst regulators (AP2III-4, AP2III-1 and AP2XII-3 [45]). Interestingly one of the most significantly upregulated genes, *srs22a* (TGME49_238440), was recently proposed as an *in* vivo-specific surface protein that demarks a subset of bradyzoites (Supplementary Figure 40)[46].

The number of differentially expressed genes between *in vitro-* and *in* vivo-derived bradyzoites was too large (n = 2,107 Supplementary Figure 4C) to guide the search for BCC regulators. We therefore reasoned that the decision point to remain arrested in G1 or progress through either cell cycle (UCC/BCC) would be found within G1 and we established a pipeline (Figure 3E) to identify drivers of BCC replication. We first examined if *in* vitro-derived tachyzoites and bradyzoites were divergent within their G1 cell cycle state. In line with conversion from tachyzoites to bradyzoites, differential gene expression analysis (Supplementary Figure 4E and Supplementary Table 3) identified 54 upregulated genes (log2FC >2) including bradyzoite specific markers and CWPs and downregulation (log2FC < −2) of 59 tachyzoite expressed genes including *sag1* and *enolase2.* As neither *in* vitro-derived tachyzoites nor bradyzoites utilize the BCC, these genes were not considered drivers of entry and were excluded from our prioritization.

As visualization of *in vitro* and *in vivo* G1 states (Figure 3F) showed little to no overlap in their clustering we performed differential gene expression analysis between these populations (Supplementary Figure 4F and Supplementary Table 4). We identify 676 upregulated (log2FC > 2) and 1146 downregulated (log2FC > −2) genes *in vivo.* As this did not sufficiently reduce the number of differentially expressed genes, we prioritized candidates that are specifically expressed during G1 by requiring overexpression (>2 log2FC (pval < 0.05)) when compared to S-phase and M-phase (Supplementary Table 2). This retained 41 genes that may regulate entry into the BCC (Figure 3G). Interestingly, a relatively unstudied AP2-domain containing protein TGME49_215895 (Figure 3H), now called AP2XI-6 according to standard nomenclature [32] was amongst the G1-specific expressed *in vivo* upregulated genes.

### AP2XI-6 is dispensable for acute stage growth

We hypothesized that AP2XI-6 acts as a transcription factor to drive replication through the 8CC resulting in *in vivo* cyst development and maturation. If AP2XI-6’s role is to regulate entry into the 8CC we would expect it to be dispensable for UCC-dependent growth, including tachyzoite growth and *in* vitro-induced differentiation. To test this, we generated a knockout in a parental line (87) that robustly forms cysts *in vitro* and *in vivo. AP2XI-6* knockout was confirmed by PCR (Figure 4A and B). As would be predicted by its fitness score (0.74, [47]), we observed no reduction in the number (Figure 4C and D) or size (Figure 4C and E) of plaques formed between the parental (87) or knockout (ΔAP2XI-6) tachyzoites. In mice infected with an inoculum size predicted to be 30-50% lethal without prophylaxis, we observed no difference in virulence based on mortality or weight loss (Figure 4F and G). Taken together, these data demonstrate that UCC-driven acute stage growth is unaffected by loss of AP2XI-6 expression.

**Figure 4.**
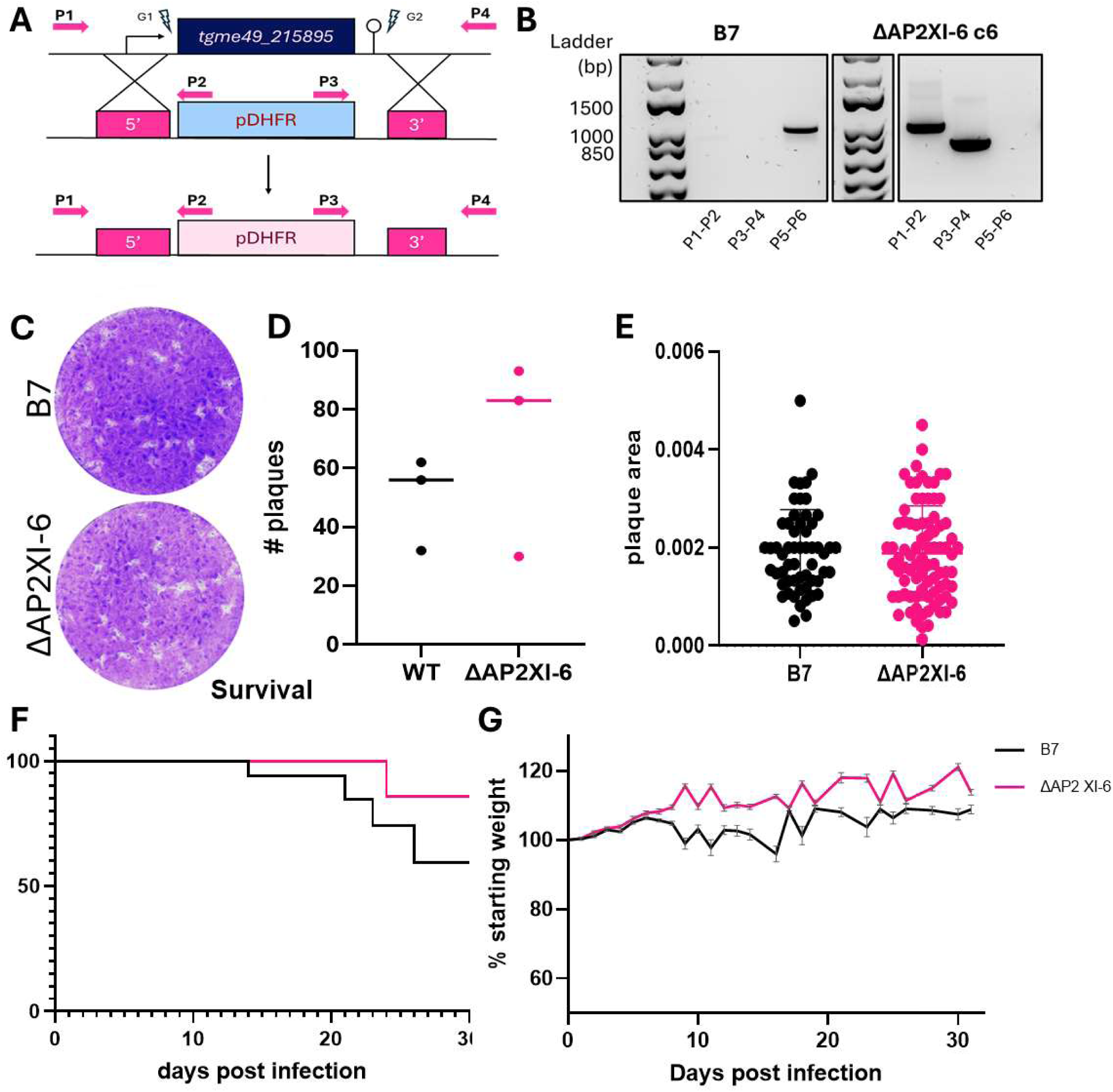
AP2XI-6 is dispensable for acute stage growth. **(A)** To generate an AP2XI-6 knockout the coding sequence was replaced with the pDHFR selection cassette that confers resistance to pyrimethamine. **(B)** Replacement of the gene with the selection cassette was confirmed by integration PCR, primer locations shown in (A). Loss of AP2XI-6 expression does not alter *in vitro* tachyzoite growth, as demonstrated by no significant difference in **(C)** plaque assays in the **(D)** number or **(E)** area of plaques formed, Confirmed by Students T-test. **(F)** Infection of mice (equal number male and female mice, no prophylaxis) with 100 B7/WT (black line) or ΔAP2XI-6 (pink line) parasites shows no significant difference in acute stage survival or **(G)** weight loss through the acute stage. Significance calculated using a Kaplan Maeir test.

### AP2XI-6 is required for chronic infection *in vivo*, but not *in vitro*

To determine if AP2XI-6 plays a role in chronic infection, mice were infected with a sublethal dose of 87 or ΔAP2XI-6 parasites and a chronic infection allowed to develop. As previously noted, we saw no difference in acute stage virulence or weight loss (Supplementary Figure 5A). We then quantified the number and size of cysts based on Dolichos lectin staining of the cyst wall. Representative cysts are shown in Figure 5A. We detected cysts in only 3/40 mice infected with ΔAP2XI-6 tachyzoites and the number of cysts per brain was significantly lower that 87 (Figure 58) with an average of 6.5 and 821.7 respectively. The average cyst size between 87 and ΔAP2XI-6 cysts was not statistically significantly different, with averages of 21.9µm and 19.8µm respectively. In line with the known sex specific differences in infection we observed more and larger cysts in female mice (average 1264.3 cysts per brain and 23.8µm diameter) compared to males (average 495.6 cysts at 19.3µm) in our wild type (B7) infections (Supplementary Figure 5B).

**Figure 5.**
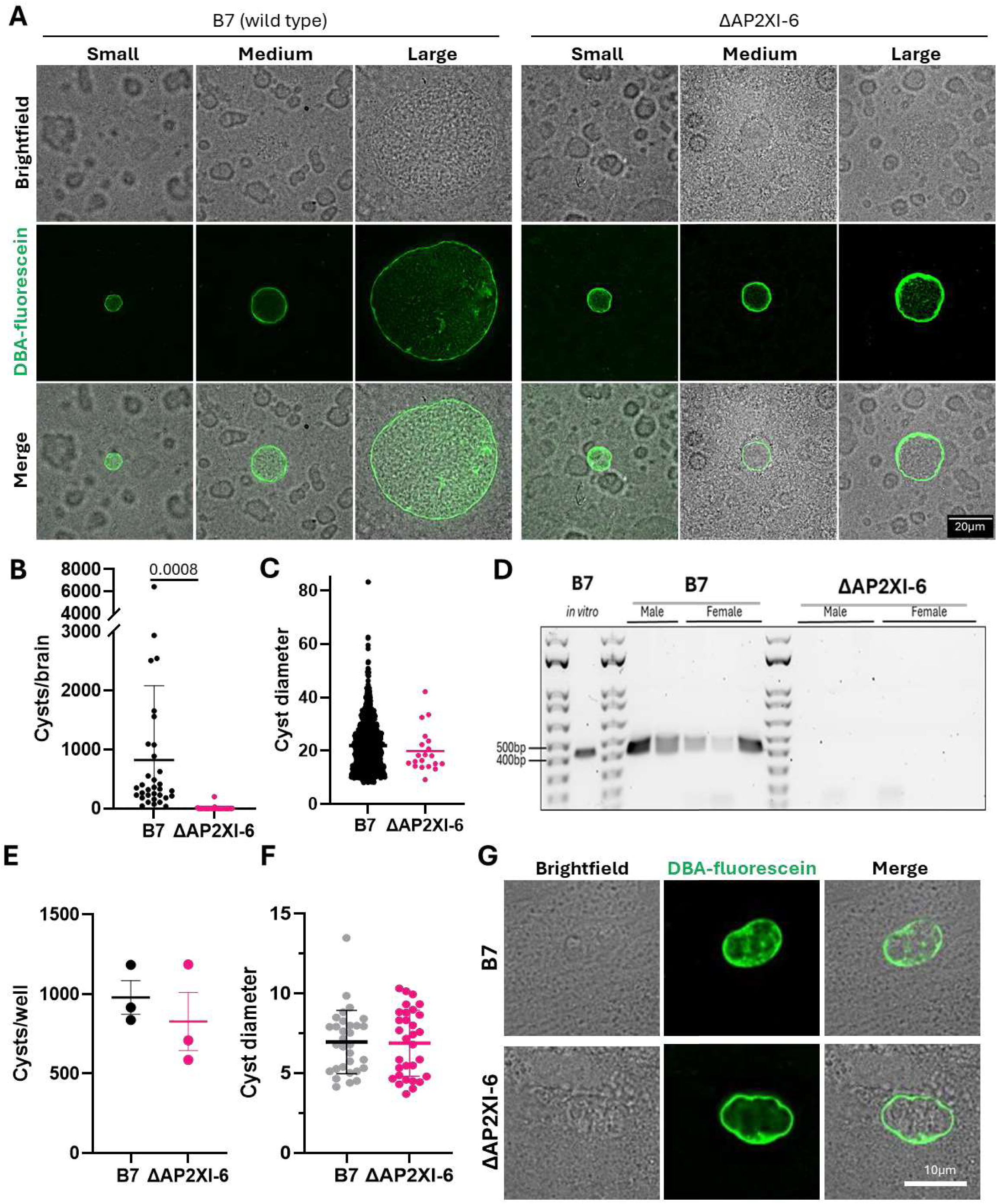
AP2XI-6 is required for an in vivo chronic infection, but not an in vitro chronic model. Following establishment of a chronic infection in vivo (30-31 days) brains were isolated and cysts stained with fluorescein-conjugated Dolichos lectin (OBA-FL). **(A)** representative examples of small, medium and large cysts from a B7 (wild type) or ΔAP2XI-6 infection. **(B)** The number of cysts per brain was quantified. The average number of cysts in a B7 infected mice was 821.7 cysts per brain while only 3 mice became infected with ΔAP2XI-6 cysts, resulting in an average burden of 6.5. **(C)** 1000 B7 and all ΔAP2XI-6 cysts (n=20) were measured at their widest point to quantify cyst size. The average size of a cyst in the wr infection is 21.91µm while ΔAP2XI-6 cysts are 19.84µm in diameter. **(D)** B1 gene PCR amplification was performed on whole DNA extracted from the brains of 5 B7 and 7 ΔAP2XI-6 challenged mice to determine if parasite DNA was present 30-31 days post infection. **(E-G)** Monolayers were challenged with equal numbers of B7 and MP2Xl-6 tachyzoites before inducing differentiation for 5 days. The number **(E)** and diameter (n=30) **(F)** of the cysts were quantified based on OBA-FL staining. **(G)** Examples of representative cysts are shown. Significance was calculated using Students t-test for B, C, E and F. Only significance< 0.05 displayed.

To determine if an MP2Xl-6 infection in the surviving mice persists in the form of pseudocysts (that do not stain with OBA) or acute stage parasites, we performed PCR for the multicopy B1 on DNA isolated from the brains of B7 and MP2Xl-6 infected mice from one biological replicate of infections (Figure 5B and C). We see amplification of the parasites B1 gene from all B7 infected mice, while none of the ΔAP2XI-6 infected mice yielded a detectable gene product (Figure 5D) demonstrating that ΔAP2XI-6 infection is cleared.

As AP2XI-6 expression is largely restricted to *in vivo* G1a we wondered if loss of expression would also alter encystation following *in vitro* differentiation. 5-days following stress-induced differentiation we quantified the number (Figure 5E) and size (Figure 5F) of cysts per well and see no difference. The cysts also appear similar morphologically (Figure 5G).

Taken together, these data show that AP2XI-6 is required for the development of a viable chronic *T gondii* infection *in vivo,* where parasites predominantly replicate through the bradyzoite cell cycle (BCC) but is dispensable for the development of a chronic infection *in vivo,* where replication through the universal cell cycle (UCC) or G1 arrest occurs.

## Discussion

To maintain a chronic *Toxoplasma* infection tachyzoites convert to bradyzoites and encyst in key host tissues including the brain. To understand how bradyzoites maintain this infection we performed scRNAseq on parasites isolated from the brains of chronically infected mice. Unlike tachyzoites, which ubiquitously express canonical stage-specific markers, bradyzoites exhibit heterogeneity in expression of known stage-specific markers. While some of these markers (e.g. *ldh2)* are variably expressed across all populations with high penetrance, others like *bag1* and *enolase1* are lowly expressed by only a few individuals in some subpopulations but highly with high penetrance in others (Figure 1E). This, along with recent data [28, 46], suggests *in vivo* bradyzoites exist within distinct subpopulations.

Variable gene expression in different subsets of bradyzoites also occurs with cyst wall proteins. This led us to hypothesize that CWP expression is spatially regulated, with expression confined to parasites at the periphery of the cyst for more direct protein delivery, utilizing the previously identified vesicular network [13]. However, using RNAscope we saw no differences in CWP transcript abundance across the cyst, demonstrating that the subpopulation of bradyzoites that express CWPs are not spatially regulated (Figure 2). Co-expression analysis revealed that CWP expression is regulated in a cell cycle dependent manner, with G1b accounting for CWP expression (Figure 2H). We propose that this cell cycle regulation ensures maintenance of the cyst wall without continual production of CWPs that may result in a thick or dysfunctional cyst wall. This hypothesis is supported by work showing that while many CWPs are required for chronic infections [11, 16, 17], a disordered and thickened cyst wall results in clearance by the host [48]. In addition, we observe dynamic cyst behavior *in vivo* where progeny cysts appear to bud from a parent cyst (Supplementary Figure 3) which likely requires a dynamic cyst wall. Changes in the rate of cyst fission have been recently reported both *in vitro* and *in vivo* following genetic manipulation to deplete the chitinase-like protein CLP1 [48] and the bradyzoite-specific cyclin, CYC5 [49].

As only 35% of the parasites express CWPs and G1b transcripts, we assigned cell cycle state to all parasites (Figure 3A). This revealed the presence of duplicated S-phase, M-phase and cytokinesis clusters that branch from a single G1a/b cluster. Integration of existing scRNAseq from *in* vitro-derived tachyzoites (Figure 3B) and bradyzoites (Figure 3C) [28] revealed that one of these loops aligns with the known *Toxoplasma* cell cycle, and we refer to this as the universal cell cycle (UCC). The second loop is poorly represented by *in* vitro-derived parasites, so we designate this as the bradyzoite cell cycle (BCC). This analysis revealed that unlike *in* vitro-derived bradyzoites, where the majority (>90%) are annotated as G1 [28], *in* vivo-derived bradyzoites have a lower proportion of G1 “arrested” parasites (∼65%) and the remaining parasites can replicate either through the UCC (∼10%) or the BCC (∼25%), highlighting a limitation in the current *in vitro* stress-induction model. Several new *in vitro* differentiation models that use overexpression of differentiation-driving transcription factors [5, 6] or cells that induce spontaneous differentiation [26, 50] have recently been developed and may offer more complete representation of *in vivo* bradyzoites. Further work will be needed to characterize bradyzoites derived from these methods. Additionally, our analysis was performed on chronic infections that had been established for 32days; it would be interesting to correlate the proportion of parasites replicating through each cell cycle (UCC/BCC) during the ∼8 week cyclic pattern of increasing and decreasing replication frequency documented when visualizing parasite replication using microscopy [20, 21, 29].

In addition to identifying two distinct cell cycle loops traversed by *in* vivo-derived bradyzoites, we identified a transcription factor, *AP2XI-6* that is only expressed by *in vivo-*derived bradyzoites during G1 (Figure 3G and H). We generated a knockout of this gene in a cystogenic parasite strain and determined that UCC-driven processes, including tachyzoite growth and *in vitro* encystation (Figure 4 and 5 E-G), are unaffected. However, BCC-dependent growth, determined by *in vivo* cyst formation, is significantly impaired (Figure 5). Taken together these data suggest that AP2XI-6 acts as a regulator of bradyzoite replication through the BCC and in its absence replication can only progress through the UCC which is not sufficient to maintain a chronic infection *in vivo*.

We propose that *Toxoplasma* utilizes these two cell cycles to ensure persistence of the chronic infection. In this model (Figure 6), all parasites that pass through G1b express CWPs to maintain the cyst wall, regardless of the cell cycle they will utilize for replication. This mechanism may have developed to replenish essential cyst wall proteins that shield bradyzoites from immune recognition while preventing excessive cyst wall growth that may impede parasite replication, cyst growth [29], cyst dynamics (as seen in Supplementary Figure 3) and cyst turnover [3, 48].

**Figure 6.**
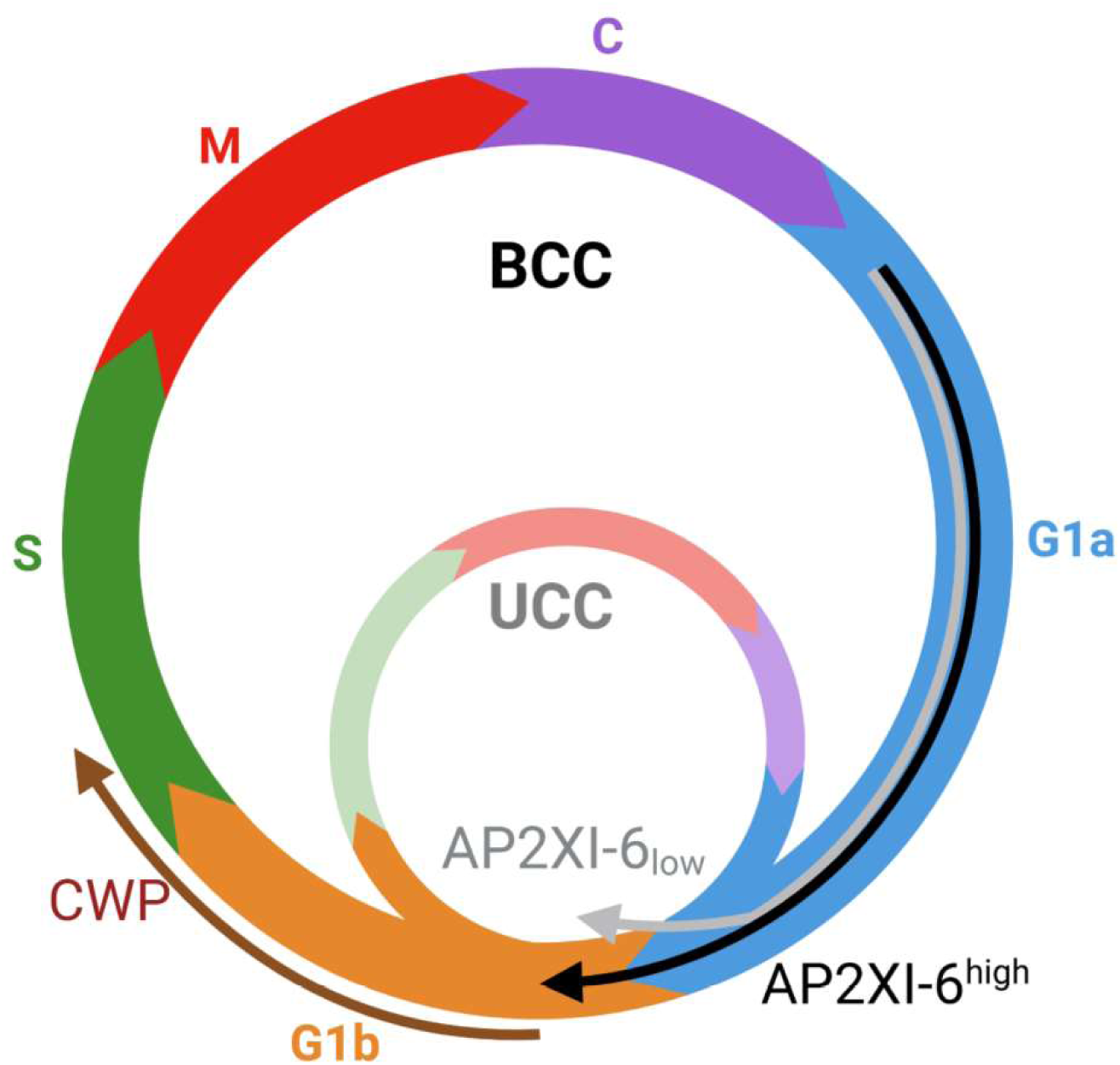
Model for the role of concurrent cell cycles in vivo. Bradyzoites in the G1b cell cycle stage (orange) express cyst wall proteins allowing replenishment of critical CWPs. Following exit from G1 in vivo bradyzoites can replicate through the canonical or universal cell cycle (UCC) or through the newly identified in vivo bradyzoite-specific cell cycle (BCC). The decision point for which cell cycle to utilize occurs in G1a (blue) with expression of AP2XI-6. High expression of AP2XI-6 initiates a cascade that drives replication through the BCC; in the absence of AP2XI-6, replication progresses through the UCC.

To balance cyst reseeding and cyst growth we propose that the UCC enables rapid parasite replication, likely required following re-seeding of cysts or cyst budding events, and replication through the BCC results in slow within-cyst replication [29]. Acting as the gatekeeper of these cell cycles, AP2XI-6 would drive replication through the BCC and in its absence, replication would default to the UCC. Balancing these cell cycles may be necessary to ensure cyst replenishment as the infection persists without causing damage to the host that may alert the immune system and result in clearance.

### Parasite culture

ME49-mCherry, B7 and ΔAP2XI-6 tachyzoites were propagated in human foreskin fibroblasts (HFFs, ATCC). These cells were grown to confluence in D10 (Dulbecco’s Modified Eagle’s Medium (DMEM) (Gibco) containing 10% v/v heat-inactivated fetal bovine serum (FBS) (Life Technologies, Carlsbad, CA), 10 mM HEPES pH 7, and 100units/ml penicillin and 100ug/ml streptomycin), as previously described [27]. Once cells reached confluence, the medium was changed to D3 (DMEM supplemented with 0mM HEPES pH 7, 100U/ml penicillin, 100ug/ml streptomycin, and 3% FBS) for the remainder of the infection with T. gondii. ME49-mCherry parasites used early in the project for scRNAseq were cultured in HFFs provided by Dr. Thomas Moehring, and cultured as above except that infections were maintained in D1 (DMEM as above containing 1% v/v FBS).

Differentiation to bradyzoite-stage parasites in vitro was induced by replacing the media 2-4hours after challenge with tachyzoites to differentiation medium (RPMI supplemented with 10mM HEPES pH 8.1, 100units/ml penicillin and 100pg/ml streptomycin, and 3% viv FBS) and grown in ambient CO_2_ conditions, as previously described [3, 27]. For all in vivo bradyzoite assays the differentiation medium was replaced daily for 5 days.

### *in vivo* infections

Infections were carried out in 4—5-week-old female (for ME49 mCherry parasites) or 6-8-week-old female and male (for B7 and ΔAP2XI-6 parasites) CBA/J mice purchased from Jackson Laboratories (Bar Harbor, ME). All mice were socially housed and acclimated for at least 3days prior to infections. Daily animal health monitoring began the day of infection. Mice were euthanized if predetermined thresholds were exceeded according to predefined welfare standards.

### Single cell RNAseq

Thirty-two days post infection with ME49 mCherry tachyzoites, mice were euthanized with CO2 and brains were isolated. Brains were washed in 5ml PBS before half of each brain was fixed in 4% paraformaldehyde and half was pooled (12mice total) before homogenization in PBS (2ml/brain) using a Dounce homogenizer. Homogenized tissue was serially passed through 18G, 20G and 22G needles five times, and centrifuged at 2000xg for 10minutes. Each sample was resuspended in 6ml 20% dextran-150 in PBS and centrifuged at 400xg for 10minutes. The myelin layer and dextran-150 were removed by aspiration. The pellet was washed once with 10ml PBS by centrifugation at 2000xg for 10 minutes. To release the bradyzoites, the pellet was resuspended in 6ml acid pepsin (AP; stock solution of [0.01 mg/ml pepsin in 1% NaCl pH 2.1] diluted in 6ml freshly made 1% NaCl pH 2.1 immediately before use) and incubated at 37°C for 5minutes. The suspension was then passed through a 26G needle directly into 12ml Na2CO3 to inactivate the pepsin [24, 25]. Released parasites were resuspended in FACS buffer (1x PBS with 1% FBS) before sorting with a BD FACSAria to isolate a pure population of viable parasites. 20,000 “Live” (DAPI negative) parasites (mCherry positive) were sorted at 4°C into D10. These parasites were concentrated by centrifugation at 2000xg for 5minutes before being resuspended in 10µl PBS. 1µl parasites were added to 9µl PBS and concentration was quantified using a hemacytometer. The solution was then diluted to 1200 parasites/µl in PBS. Cell partitioning was completed using the Chromium Next GEM Single Cell 3’ GEM Kit v3.1 following manufacturer’s instructions [37]. Cells were combined with barcoded Single Cell 3’ v3.1 Gel Beads, a Master Mix containing cells, and Partitioning Oil onto Chromium Next GEM Chip G to create GEMS. Cells were loaded according to the manufacturer’s instructions to recover 8000 cells. Barcoded full-length cDNA from GEMS is then pooled and subjected to PCR amplification (13 cycles) to generate sufficient mass for library construction. Libraries were prepared for sequencing using the NextSeq kits and sequenced on a NextSeq2000 at a read depth of 50,000 reads per cell, using paired-end sequencing (150bp).

### Single-Cell RNA Sequencing and Data Processing

Sequencing reads were aligned to the *Toxoplasma gondii* ME49 reference genome (annotation version 65) using Cell Ranger v7.2.0 (1Ox Genomics). Quality control of the resulting raw count matrix retained 6,505 cells for downstream analysis. Counts were normalized by library size, log1p-transformed, and scaled to unit variance across cells. Principal component analysis (PCA) was performed, and a k-nearest neighbor graph was constructed using the cosine distance metric. Community detection using the Leiden algorithm identified six transcriptionally distinct clusters.

### Cross-Dataset Label Transfer

Cell type and phase labels were transferred across datasets using two-set canonical correlation analysis (CCA) integration [51] combined with batch-balanced k-nearest neighbors **(BBKNN)[52].** Each query cell was matched to its 19 nearest neighbors in the reference dataset, and the majority label was assigned. Cell cycle annotations were transferred from [28](Accession: GSM4306214), where phases had been inferred from DNA content and unsupervised clustering.

### Downstream RNA-seq Data Analysis

All single-cell analyses were performed using SCANPY [53] and SCLab [https://github.com/umbibio/sclab]. Differential gene expression was assessed with sc.tl.rank_genes_groups in SCANPY, which implements the Wilcoxon rank-sum test for pairwise comparisons between groups.

### RNAscope

#### RNAScope Multiplex Fluorescent Assay V2 on the Leica BOND Rxm

Five-micron sections of mouse brain tissue were cut onto glass slides. RNA fluorescence *in situ* hybridization (FISH) was performed using the RNAScope Multiplex Fluorescent Assay V2 on the Leica Biosystems’ BOND RXm Research Advanced Staining System. Slides were baked and dewaxed on the BOND RXm then antigen retrieval was performed using BOND ER2 solution for 15minutes at 95°C. They were then treated with RNAscope® 2.5 LS Protease Ill from ACD for 15minutes and RNAscope® 2.5 LS Hydrogen Peroxide for 10minutes, both at 44°C. The sections were hybridized with either a test probe cocktail consisting of C1-gondii-SRS44/C2-gondii-TGME49/C3-LDH2, the mouse RNAScope® 2.5 LS Multiplex Positive Control Probe (C1-POLR2A, C2-PPIB, C3-UBC) or the RNAScope® 2.5 LS Multiplex Negative Control Probe (DapB) for 2hours at 42°C. The target probe cocktail was made by diluting the C2 and C3 probes 1:50 into the C1 probe. After the 2hour hybridization, the probe signal was amplified through three 30minute incubations at 42°C with RNAscope® LS Multiplex AMP 1, RNAscope® LS Multiplex AMP 2, and RNAscope® LS Multiplex AMP 3. Fluorescent signal was developed for each channel using RNAscope® LS Multiplex HRP and Opal™ fluorophores from PerkinElmer diluted 1:500 in TSA Buffer. Opal 690 was used for the C1 channel, Opal 570 was used for the C2 channel, and Opal 520 was used for the C3 channel. Finally, DAPI was used to stain the nuclei.

#### Confocal Imaging

Stained slides were imaged on a Nikon A1R-ER point scanning confocal microscope (Nikon, Tokyo, Japan). Cysts were imaged using an Apo TIRF 60X oil DIC N2 objective (NA 1.49) with a galvanometric scanner. Image scan zoom was set to 2.0. The DAPI, Opal 520, Opal 570, and Opal 690 dyes were captured using 405nm, 488nm, 561nm and 640nm lasers and collected with 450/50, 525/50, 600/50, and 685/70 bandpass filters respectively. Laser power and gain settings were established by imaging an unstained control. Images were saved in ND2 file format in NIS Elements (version 5.2.1, Nikon, Tokyo, Japan).

### Image Analysis

Colocalization of the RNA probes was measured using Volocity version 6.3.0 (Perkin Elmer, Waltham, MA, USA). A scatterplot of fluorescent intensity for each channel was used to create thresholds for each channel by comparing positive images to the negative control image.

The images were analyzed using lndica Labs’ HALO software (v.4.0.5107.313) (Albuquerque, New Mexico USA).

Each cyst was circled, and three concentric circles were generated to designate an inner, middle, and outer layer of the cyst. The intensity of each fluorescent signal as well as the area of each fluorescent signal in each layer of the cyst was measured using the Area Quantification FL v2.3.9 algorithm. The number of RNA copies in each layer of the cyst were counted using the FISH v3.3.1 algorithm.

### lmmunohistochemistry

Brain tissue was fixed in 10% formalin, embedded in paraffin and sectioned (5µm). Deparaffinized sections were hydrated with distilled water. Sections were then stained with Mayers Hematoxylin and Alcoholic-Eosin. Slides were then dehydrated with ethanol and cleared with xylene before imaging with a Leica-Aperio VERSA 8 whole slide scanner system. Image analysis was completed using the Image scope (x64) software.

### Generation of ΔAP2XI-6 line

All synthetic oligonucleotides were synthesized by Integrated DNA Technologies and are listed in Supplementary Table 1. All sequence and gene information in this work was downloaded from ToxoDB.org [54, 55]. Design of sgRNA’s for AP2XI-6 was performed using the Eukaryotic Pathogen CRISPR guide RNA/DNA design tool.

#### Plasmid generation

The AP2XI-6 knockout plasmid and two sgRNA CRISPR plasmids were generated by Gibson assembly. For the knockout plasmid (pΔAP2XI-6), the pDHFR floxed vector (synthesized by Twist Bioscience) was linearized by digestion with Aflll (NEB), once fully linearized this was combined with a gene fragment containing 200bp 5’ and 3’ homology to the AP2XI-6 promoter and 3’UTR (synthesized by Twist Bioscience) using the Gibson Assembly Master Mix kit (NEB) following manufacturer’s instructions. As per manufacturer’s instructions 5µl of the reaction was transformed into dh5a bacteria (NEB) before recovery in SOC media and selection on ampicillin plates (100mg/L). Individual colonies were grown overnight before miniprep (Promega Wizard miniprep kit) and screening for insertion of the homology arms by restriction digest and sequencing to confirm integration (OMRF Clinical Genomic Center). For the two sgRNA CRISPR/cas9 plasmids (sgRNA:AP2XI6 a and b), the parental plasmid, PSS013, containing the Cas9 gene was linearized with Bsa-I (NEB) and 20bp sgRNAs were integrated using the same Gibson assembly method above, insertion of the sgRNA was confirmed by plasmid sequencing (OMRF Clinical Genomic Center).

#### Transfections

To generate the knockout parasites approximately 1×10^7^ B7 tachyzoites were transfected with pΔAP2XI-6 (50µg) and sgRNA:AP2XI6 a and b (5µg) resuspended in P3 buffer using a 4D-nucleofector (Lonza) with pulse code Fl-158. Transfected parasites were selected with 3µM pyrimethamine final concentration in D3. Following drug selection clones were isolated by limiting dilution (5 parasites per well) and integration was confirmed by diagnostic PCR *(Taq* polymerase (New England Biolabs)) of isolated genomic DNA (isolated as crude lysate with 2ul Proteinase K (Thermo Scientific) and 98ul PBS (Gibco)).

### Plaque assay

Tachyzoites (B7 and ΔAP2XI-6) were harvested from large vacuoles within HFF cells before counting on a hemocytometer. Five hundred parasites were inoculated per well (6 well plate) onto freshly confluent HFF monolayers D3 and left undisturbed for 21 days in a 37°C, 5% CO2 incubator. Plaque formation was assessed by counting the number of zones of clearance on EtOH-fixed, crystal violet-stained HFF monolayers. Each stained plate was scanned with a high-definition digital scanner (Epson) to obtain images and to quantify plaque area. For plaque area measurements, all discernible plaques from all stained wells were analyzed using lmageJ to define the plaque area in square pixels (pix2). The number and size of plaques was compared between the two lines using a Students t-test.

### *in vivo* virulence

Mice were infected with 100 B7 or ΔAP2XI-6 parasites, known to result in 30-50% mortality for B7, and their weight monitored daily for the first 14 days of infection then thrice weekly until 30 days post infection. If mice reached predefined welfare standards they were euthanized.

### *in vivo* cyst quantification

Mice were euthanized and their brains isolated 30-31 days post infection. Each brain was washed with 1ml PBS before a quarter of each brain was homogenized with a Dounce homogenizer. Homogenate was passed serially through 18G, 20G needles 4x each before centrifugation at 5000xg for 8 minutes. Cysts were stained with DBA-Fluorescein, a fluorescently conjugated lectin that stains the cyst wall (Vectorlabs). Cysts were fixed (4% PFA, 1ml/quarter brain) for 15 minutes before permeabilization and blocking (1% BSA, 0.4% Triton x-100 in PBS 1ml/quarter brain) for 1 hour. The cyst wall was then stained with DBA-Fluorescein (1:100 1% BSA in PBS, 100µl/quarter brain) for 1 hour in the dark. The stained sample was washed 2x 1ml 1% BSA in PBS and 1x PBS before resuspension in 80µl PBS. Ten microliters was applied to a slide and allowed to dry in the dark for 30 minutes before application of one drop of prolong gold antifade (lnvitrogen) and addition of a 1.0 glass cover slip. The slides were cured overnight at room temperature, in the dark, before storage at 4°C.

Cysts were quantified and measured on a Leica Thunder DMi8 Microscope at 20x magnification and imaged at 100x magnification using Leica LAS X software. Three to four technical replicates were quantified for each biological replicate. Up to 1000 cysts per replicate was measured. Cyst diameter was measured at the widest point of the cyst at the central slice of the cyst. All quantifications were completed blind and by two individuals, mean is presented.

### Cyst burden PCR

To determine if cysts were present, below the limit of detection, or if pseudocysts or tachyzoites were sustaining an infection in mice we performed PCR on DNA isolated from half a mouse brain. DNA was isolated using New England Biolabs Monarch HMW DNA Extraction Kit for Tissue and 250ng total DNA was used for a 30 cycle PCR reaction *(Taq* polymerase (New England Biolabs)) with the B1 primers (Supplementary Table 1) [56] that amplify a product of 451 bp from the multicopy gene.

### *in vitro* cyst quantification

To determine if in the absence of AP2XI-6 cysts were formed following induction to differentiate through stress we performed *in vitro* differentiation assays. 8,000 WT or ΔAP2XI-6 tachyzoites were added to a 24 well plate containing a confluent monolayer of HFF cells. 4-hours post inoculation the media was replaced with differentiation medium (RPMI supplemented with 10mM HEPES pH 8.1, 100 units/ml penicillin and 100 µg/ml streptomycin, and 3% v/v FBS) and grown in ambient CO2 conditions, as previously described [3, 27]. Differentiation medium was replaced daily for 5 days and cells were fixed with a 2% paraformaldehyde solution for 15 minutes at room temperature. Fixed cells were blocked and permeabilized with 1% BSA, 0.4% Triton x-100 in PBS for 1 hour. The cyst wall was then stained by DBA-Fluorescein at 1:100 in 1% BSA in PBS in the dark for 1hour. Stained slides were washed twice with 1% BSA in PBS and once with PBS before the application of one drop of prolong gold antifade (lnvitrogen) and addition of a 1.0 glass cover slip. Slides were then allowed to cure overnight at room temperature, in the dark, before storage at 4°C. Cysts were quantified using a Leica Thunder DMi8 Microscope at 20x magnification and imaged at 100x using Leica LAS X software. 3-4 technical replicates were quantified for each biological replicate.

### Statistical analysis

Statistical analyses for cyst counts, cyst diameter, plaque number and plaque size were conducted in GraphPad Prism (version 10.2.3) using two-tailed t-tests. Differences in survival were analyzed by Kaplan Maier. For differential gene expression analysis, Wilcoxon rank-sum test for pairwise comparisons was utilized. A p-value <0.05 was considered statistically significant.

## Data and code availability

scRNAseq data were deposited into GEO (insert accession number here). Custom code/pipelines can be provided upon request and will be deposited of github (link here).

## Animal Care and Ethics Statement

Mice were housed in an Association for Assessment and Accreditation of Laboratory Animal Care International-approved facility at either the University of Vermont or the University of Oklahoma health sciences center. All animal studies were conducted in accordance with the US Public Health Service Policy on Humane Care and Use of Laboratory Animals, and protocols were approved by the Institutional Animal Care and Use Committee at the respective universities.

## Supporting information

Supplementary Figures (all)

## Acknowledgements

*Toxoplasma gondii* gene information and genome annotations were accessed on http://ToxoDB.org and were instrumental for this work. Thanks to Dr. K. Brown, whose feedback on the manuscript and figures was invaluable. We would like to acknowledge the microscopy imaging center (MIC, (RRID# SCR_018821)), the Harry Hood Bassett Flow Cytometry and Small Particles Detection (FCSPD) Facility and the Vermont Integrative Genomics Resource, these core facilities made this work possible. The MIC would like to acknowledge the following equipment grants; The confocal microscope was purchased with a NIC-NCRR grant (ISI0OD025030-01), the Autostainer and HALO system were supported by an NNE-CTR NIGMS CTR grant (U54GM11516) and the Versa was purchased with a UVMMV shared instrumentation award.

This study was supported by [RK/GW], Health and Human Services grant P30GM118228 (RK/GEW), [RK/EF/GH] 5P20GM134973-05 and startup funds from the University of Oklahoma Health Science, [KZ/AA] 1R01Al167570-01A1 and 1R21Al174592-01A1.

---

**Supplementary Figure 1. Bradyzoite subclusters express characteristic markers. (A)** Heatmap showing expression of all marker genes identified across all clusters and individual cells. **(B)** Expression of microneme transcripts shows two distinct patterns; cluster 2 (green bar) into lower cluster 1 (orange bar) for *mic2* and *m2ap* known to localize to one subpopulation of micronemes (Micronemes-2, green) and predominantly cluster 1 (orange bar) like *mic3* that localize to a second subpopulation of micronemes (Micronemes-1, orange). Bradyzoite-specific microneme proteins *ama2* and *ama4* show expression analogous to *mic3* indicating localization to micronemes-1 (orange). **(C)** Known cyst wall proteins show common expression only by specific subsets of bradyzoites in clusters O (blue) and 3 (red).

**Supplementary Figure 2. Cell cycle dependent expression of marker genes.** To determine if cyst wall proteins are expressed by bradyzoites during a specific cell cycle state we performed co-expression analysis with genes known to have stage-specific expression. Here we show the expression profiles of; **(A)** All genes used in this analysis and genes most highly expressed expressed during **(B)** G1a, **(C)** G1b, **(D)** S-phase, **(E)** M-phase and **(F)** cytokinesis. Data [32] and https://toxodb.org/toxo/.

**Supplementary Figure 3. *in vivo* cysts can undergo fission.** Representative images of cysts undergoing fission where one or more progeny appear to bud from the parental cyst. **(A)** Fission begins with formation of a protrusion to make the cyst more bean shaped. **(B)** This is followed by a delineation between the two **(C)** or more cysts **(D).** Finally, the space between the cysts expands until there are clearly two separate entities. **(E)** Proportion of all cysts analyzed (88, RNA scope and 66 IHC) that are classified as cysts, loose cyst, budding cysts or multi-budding cysts. **(F)** Examples of a cyst, loose cyst and budding cyst in immunohistochemistry samples stained by H&E.

**Supplementary Figure 4. in vivo and in vitro bradyzoites show divergent gene expression patterns. (A)** in vivo isolated bradyzoites cluster into 6 distinct clusters (Figure 1). **(B)** Integration of in vitro bradyzoites with this data and transfer of cluster identities revealed that in vitro bradyzoites are largely found within cluster 0 that we show in this manuscript contain G1b parasites and express cyst wall proteins. **(C)** Differential gene expression between in vitro and in vivo derived bradyzoites revealed upregulation of 664 and downregulation of 1443 genes in vivo. These included several ribosomal proteins (grey), cytochrome proteins and known stage regulating transcription factors. All genes with a log2fold change> 8 or <-7 are annotated. **(D)** One of the most upregulated in vivo transcripts is *srs22a,* a recently identified surface protein [46] that demarks a specific subset of bradyzoites in vivo but is not expressed in vitro. **(E)** The first step of our prioritization strategy was to identify genes differentially expressed between in vitro tachyzoite and bradyzoite G1 as these would not likely be regulators of the BCC. This dataset primarily identifies stage-specific genes including cyst wall proteins (blue) and stage-specific surface markers *sag1* and *sag4.* All transcripts upregulated by log2FC > 6 or < −4 are annotated. **(F)** Differential gene expression analysis between G1 of in vitro and in vivo bradyzoites identified 676 upregulated and 1146 downregulated genes in vivo. These largely overlap with genes differentially expressed between in vitro and in vivo bradyzoites (C) and therefore required additional prioritization to identify G1 specific genes.

**Supplementary Figure 5. Sex specific differences in virulence, cyst burden and cyst size.** Mice were infected with 50 B7*/WT* (black) or ΔAP2XI-6 (pink) parasites for all experiments. **(A)** Shows no significant difference in survival or weight loss through the acute stage. Significance calculated using a Kaplan Maeir test. **(B)** Following isolation of brains, cysts were stained and quantified by fluorescein conjugated Dolichos lectin (DBA-FL) staining of the cyst wall. The average number of cysts in a WT/B7 infected mouse is 1264.3 for female mice while male mice have an average of 495.6. The average number of ΔAP2XI-6 cysts is 0.9 and 11, respectively. While the average cyst size is not different between B7 and ΔAP2XI-6 cysts, B7 infected female mice have statistically significantly larger cysts (average 23.Sµm) than male mice (average 19.3µm). Statistical difference determined by Students t-test.

**Supplementary Table 1. Oligonucleotides used in this study.**

**Supplementary Table 2. Cluster markers identified in single cell RNA sequencing.**

**Supplementary Table 3. Differential gene expression analysis between *in vitro* and *in* vivo-derived bradyzoites**

**Supplementary Table 4. Identification of G1-specific *in vivo* upregulated genes of interest.** Differential gene expression between *in vitro* and *in vivo* G1. Candidates overexpressed in G1 versus S-phae. Candidates overexpressed in G1 versus M-phase.

## Notes

### Competing Interest Statement

The authors have declared no competing interest.

