## Supplementary figures and images for "Single-cell analysis of brain-derived *Toxoplasma* bradyzoites reveals a novel cell cycle regulated by AP2XI-6"

### Supplementary Figures (all)

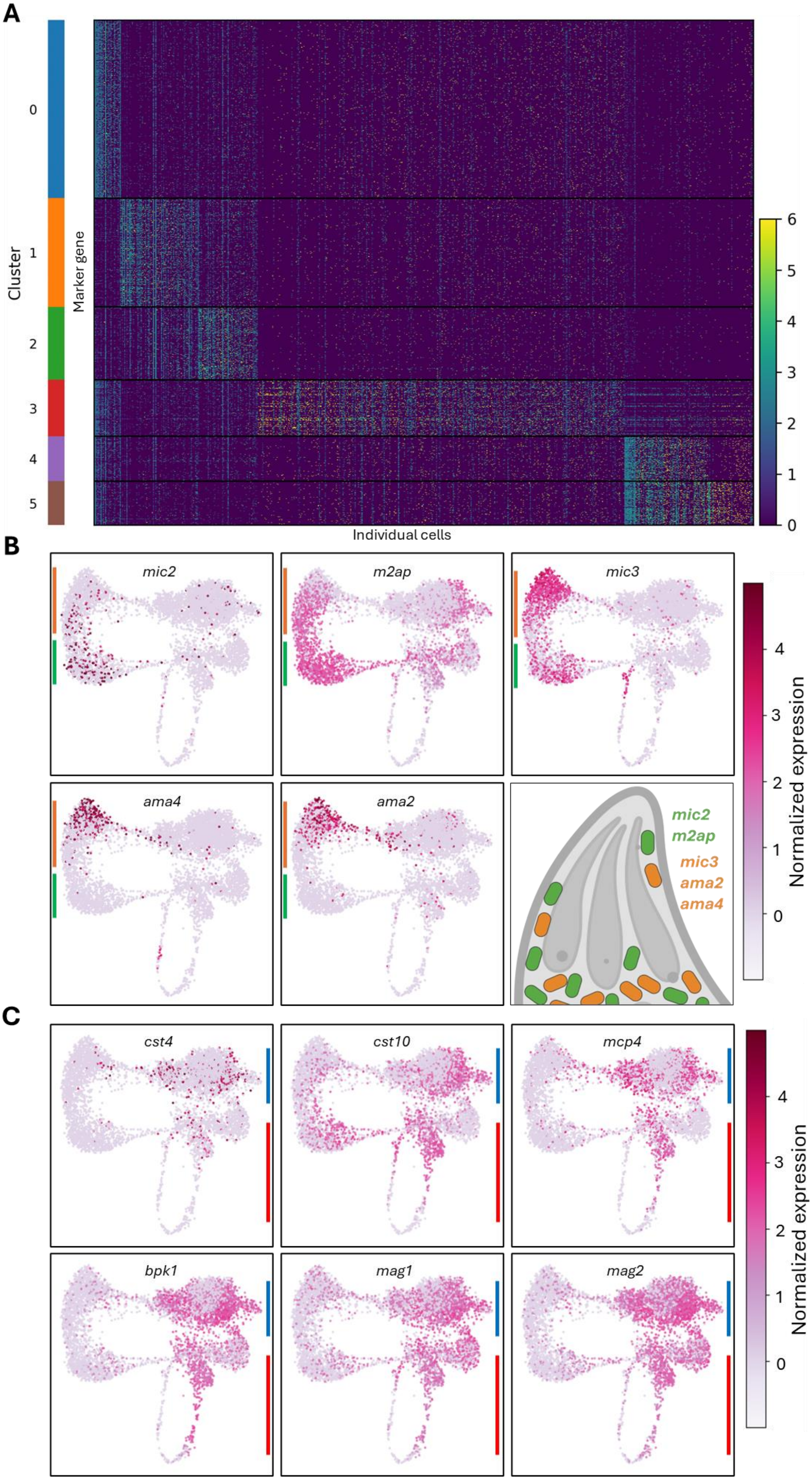

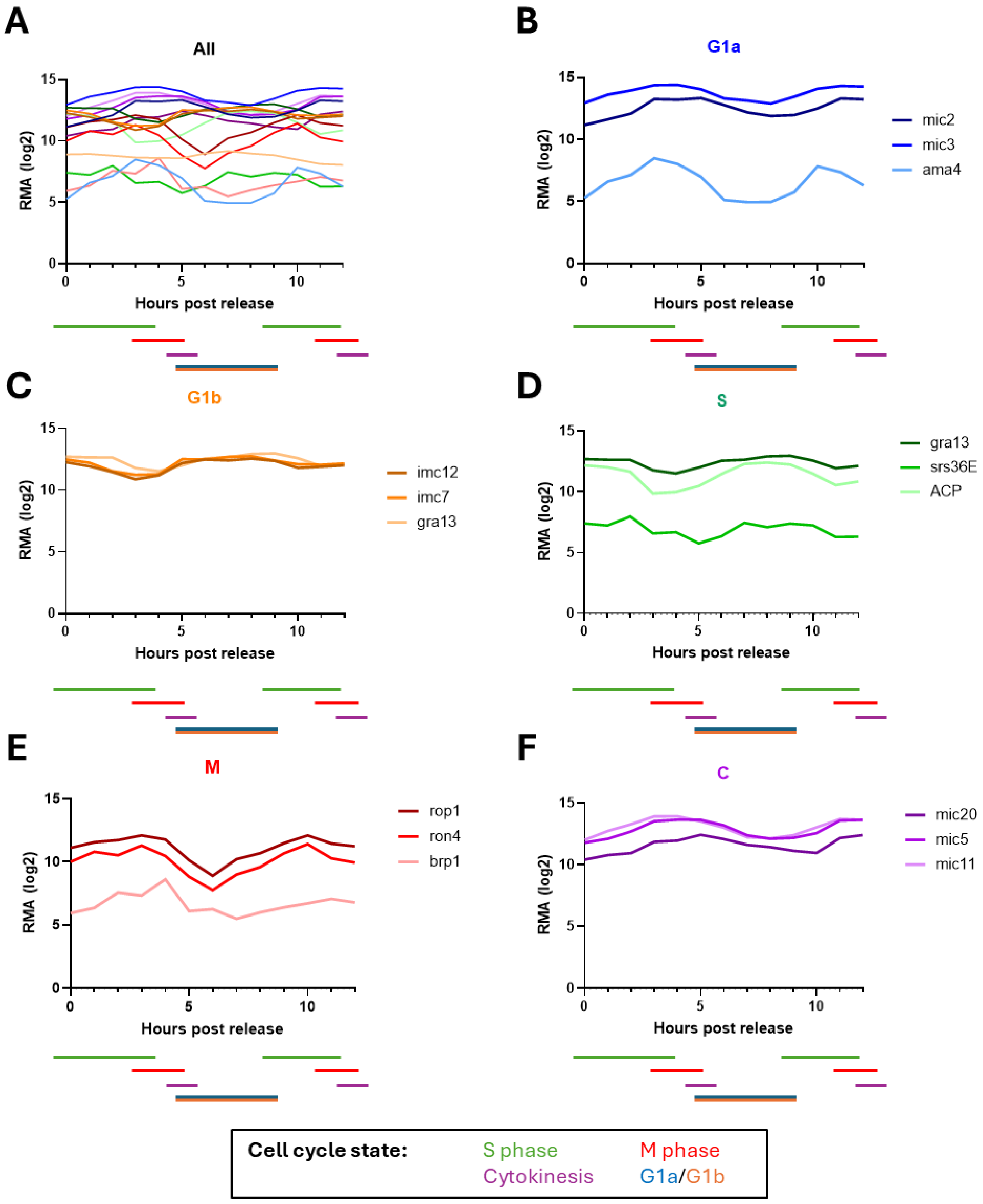

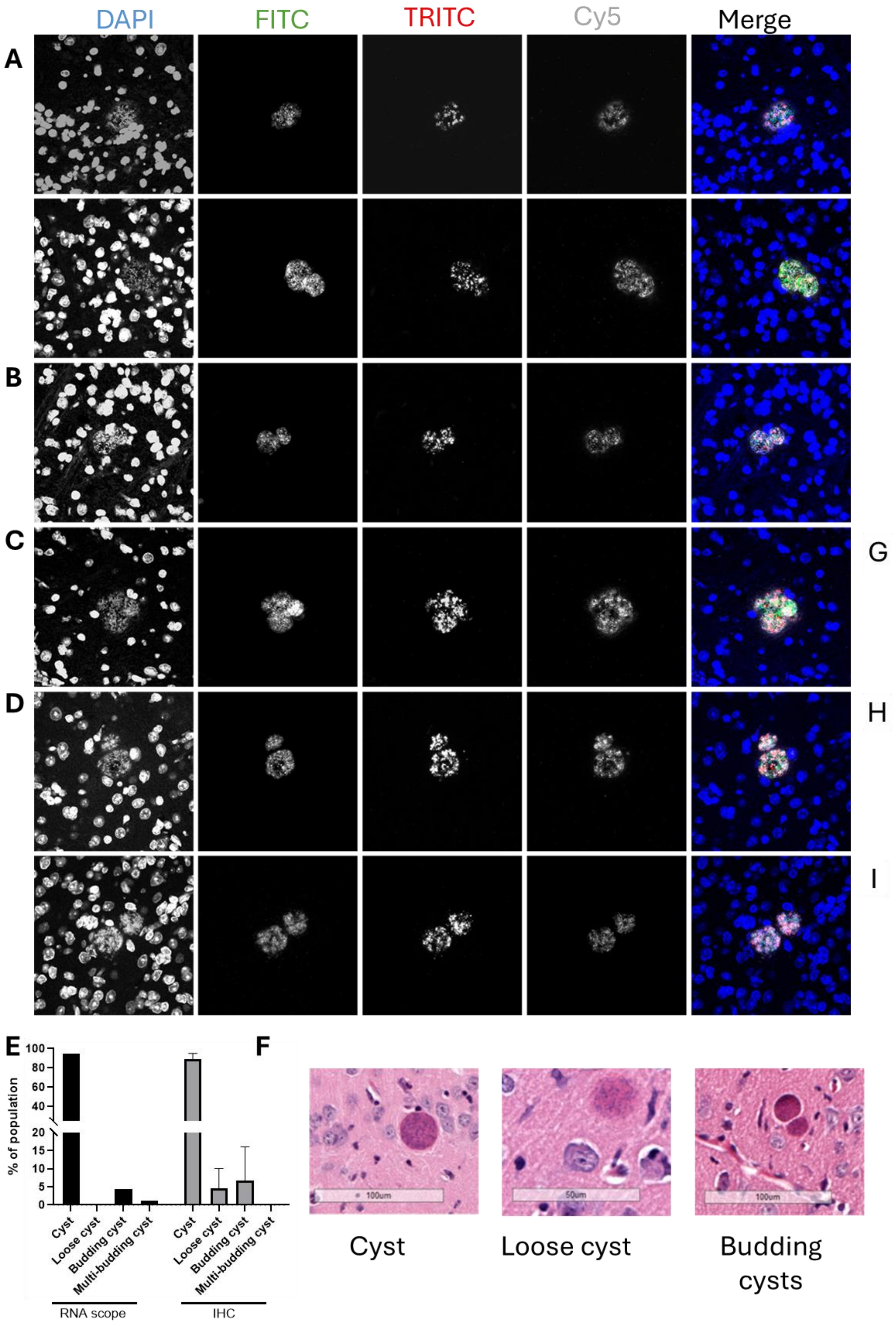

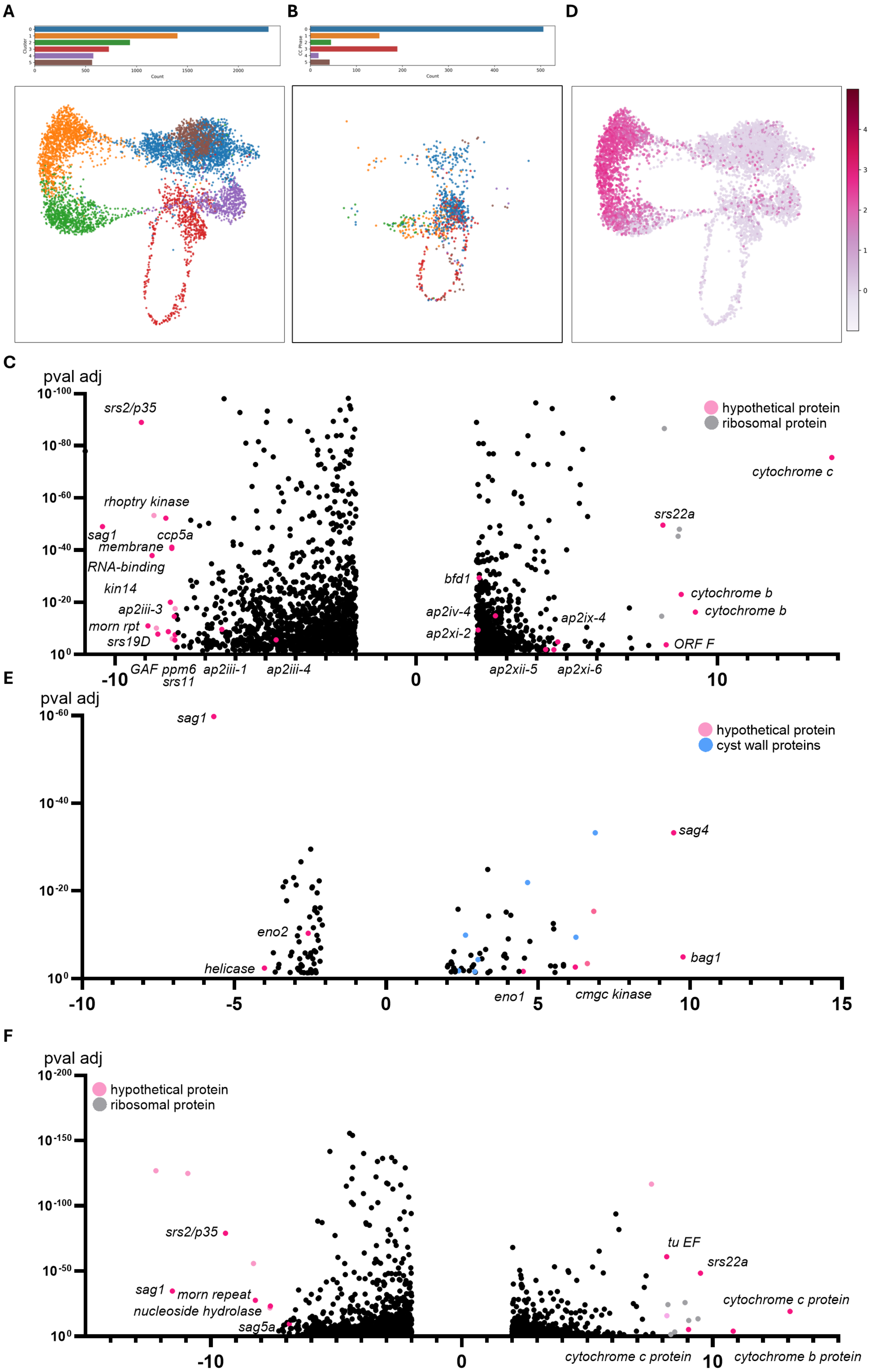

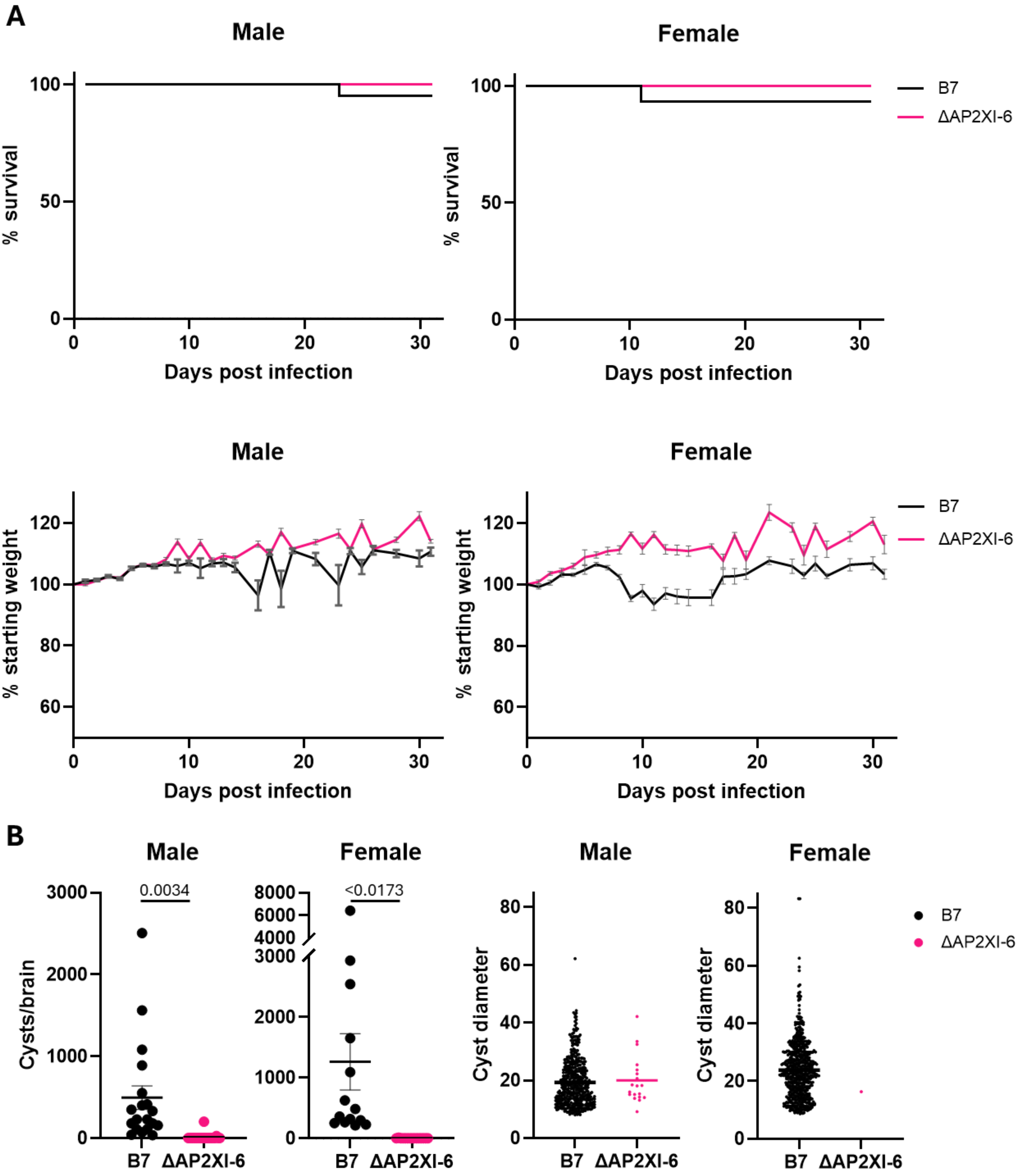
